# Predicting pyrazinamide resistance in *Mycobacterium tuberculosis* using a graph convolutional network

**DOI:** 10.1101/2025.10.28.685176

**Authors:** Dylan Dissanayake, Viktoria Brunner, Dylan Adlard, Joseph A. Morrone, Philip W. Fowler

## Abstract

**Background:** Pyrazinamide is an important first-line antibiotic for treating tuberculosis, with resistance primarily driven by mutations in the *pncA* gene. Traditional machine learning models are able to predict pyrazinamide resistance with some success, but are limited in their ability to incorporate 3-dimensional protein structural information. Graph neural networks offer the potential to integrate protein structure and residue-level features to better predict the impact of mutations on drug resistance.

**Results:** We trained a graph convolutional network on PncA variants containing missense mutations and evaluated its ability to classify resistance to pyrazinamide. Each PncA variant was represented as an amino acid-level graph, with edges calculated from 3-dimensional spatial proximity, and node features derived from chemical properties and mutation meta-predictors. We used AlphaFold2 to generate predicted structures of the PncA variants, which we used to create the protein graphs. The predicted structures of resistant PncA variants showed greater deviation from the wild-type structure compared to susceptible variants. Our model achieved an F1 score of 81.6 %, sensitivity of 81.6 % and specificity of 80.4 % on the test set and either matched or exceeded the performance of a published set of traditional machine learning models. We show that both structural graph connectivity and node features contribute significantly to model performance. Furthermore, we employ additional train/test dataset splits to demonstrate the GCN’s ability to generalise and predict resistance in samples with mutations in unseen positions and structural regions.

**Conclusions:** Our study demonstrates that graph-based deep learning can leverage protein structure and biochemical features to accurately predict antimicrobial resistance, despite being trained on a small dataset with little variation. We present this as a proof-of-concept for these methods to be applied to resistance phenotype prediction in more genetically diverse pathogens to predict the more complex observed patterns of antimicrobial resistance.

## Background

Antimicrobial resistance (AMR), particularly in the context of bacterial infections, is a growing challenge and one of the most serious threats to global public health [1– 3]. *Mycobacterium tuberculosis (M. tuberculosis)* has long been recognised by the World Health Organization (WHO) as a bacterial pathogen that is a significant concern for global healthcare and is prone to developing resistance to major antibiotics [4–7].

Tuberculosis (TB) remains the world’s leading cause of death by a single infectious agent [8]. Global TB efforts are hindered by the spread of multidrug-resistant TB (MDR-TB) strains, which are resistant to the first-line drugs rifampicin and isoniazid, and extensively drug-resistant TB (XDR-TB) strains, which are also resistant to the fluoroquinolones and at least one other second-line drug [6]. TB is treated with a regimen of multiple (usually four) antibiotics and all need to be effective to maximise the probability of a positive patient outcome. Hence, the rising proportion of strains resistant to one or more antibiotics is a significant challenge [9].

Drug resistance in TB is generally acquired by mutations occurring in chromosomal genes [6]. Exposure to TB drugs exerts a selection pressure that promotes the emergence of resistant *M. tuberculosis* strains via spontaneous mutations. These strains are associated with poor treatment outcome and often infect other people. An example of this is the acquisition of resistance to the first-line drug pyrazinamide through mutations in *pncA* [10–13]. Pyrazinamide is a prodrug that is converted to pyrazinoic acid, its active form, by the pyrazinamidase PncA, encoded by the *pncA* gene. Once converted to pyrazinoic acid, it diffuses out of the cell and disrupts membrane transport by depleting the membrane potential [14].

Mutations in *rpsA, panD* and *clpC1*, as well as in the efflux pumps *Rv0191, Rv3756c, Rv3008*, and *Rv1667c*, have also been associated with pyrazinamide resistance but these are rare [15–18]. Over 90% of pyrazinamide-resistant isolates are found to have mutations in *pncA* or its promoter region [19]. In addition to this, mutations are frequently observed to be distributed across the entire *pncA* gene (Fig. 1) and its upstream promoter, as the gene is non-essential and mutations or loss of function results in little fitness cost to the bacteria [20–22]. This is in contrast to genes such as *rpoB* and *katG* which have well-defined so-called “resistance-determining regions”, where the majority of resistance-conferring mutations to rifampicin and isoniazid, respectively, occur [23–25].

**Fig. 1:**
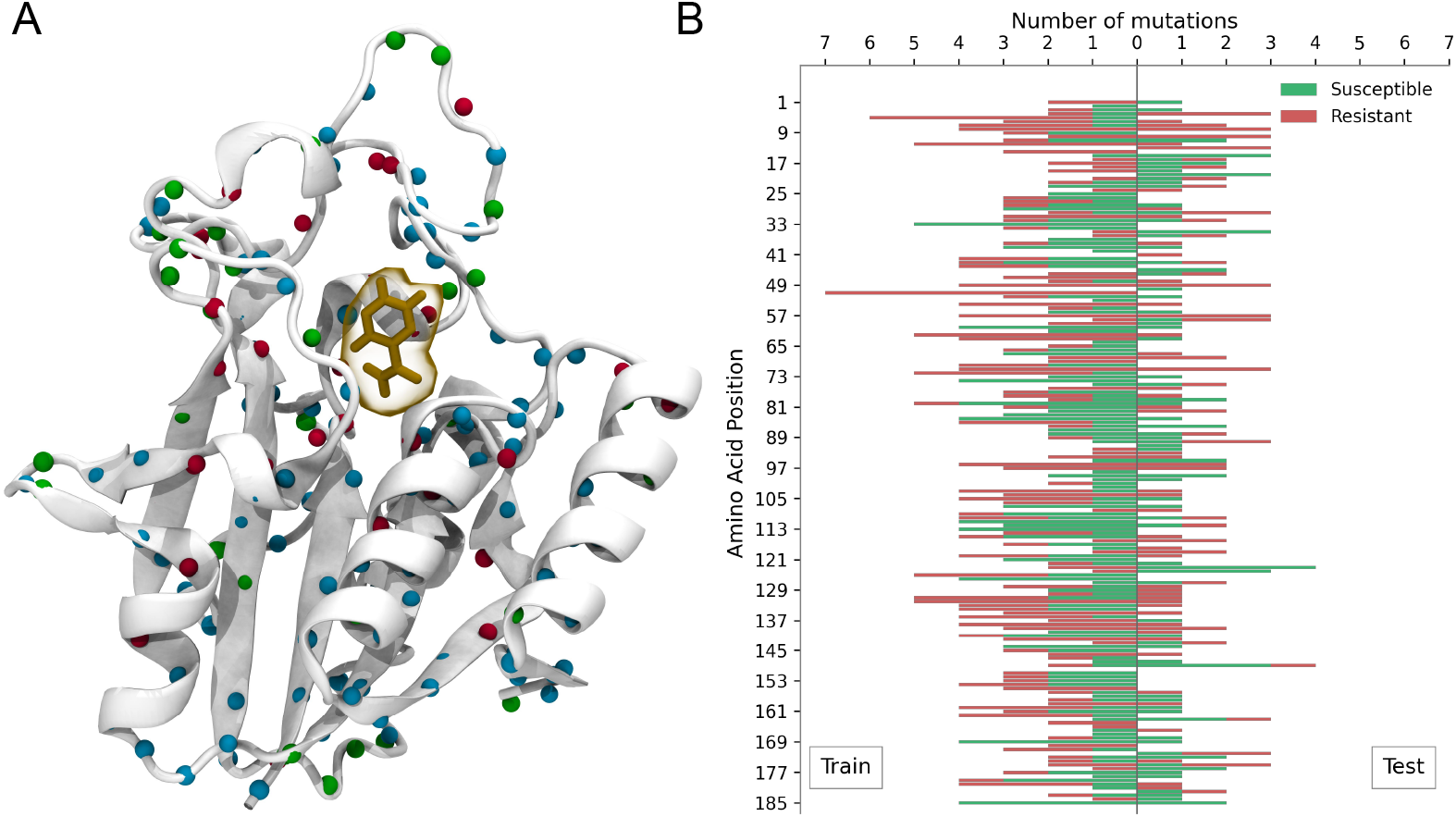
Both resistant and susceptible mutations are found throughout the full length of PncA. **(A)** PncA structure with pyrazinamide bound. Positions of mutations from the aggregated datasets are mapped onto the protein structure. Susceptible mutation positions are shown in green, resistant mutation positions are shown in red, and positions where both resistant and susceptible mutations are found are shown in blue. Bound pyrazinamide is shown in orange and the Fe^2+^ ion in purple. Structure rendered using VMD [26]. **(B)** Distribution of mutations across the PncA protein sequence. Green and red indicate susceptible and resistant mutations respectively. Bars to the left and right indicate mutations found in samples in the train and test sets respectively, as per the dataset split from Carter *et al*. [27]. Only one residue (G150) in the PncA sequence did not feature any mutations in the dataset.

To date, methods for predicting resistance from *M. tuberculosis* whole genome sequencing data rely on catalogues listing mutations that have been statistically associated with resistance to one or more drugs [28, 29]. This has been made possible by the availability of large clinical datasets where each sample has been both whole genome sequenced and undergone drug susceptibility testing (DST) [30]. This approach is fundamentally *inferential* and therefore cannot predict the effect of novel or rare mutations, as it is limited by the drugs that have been tested and the mutations observed in the samples. To tackle this issue, machine learning methods, such as logistic regression, random forests, decision trees and gradient-boosted decision trees, as well as deep learning approaches using neural networks, have all been applied to the problem of *de novo* resistance prediction in *M. tuberculosis* [31, 32]. Traditional tabular machine learning methods, however, have limitations: they are restricted to using solely genetic sequence information or features of a single amino acid mutation for their predictions, and struggle to incorporate structural information and more expressive features based on the chemical properties of individual amino acids.

Previous studies have shown that using both sequence-based and structure-based features with traditional machine learning methods can improve sensitivity and specificity of resistance prediction [12, 27]. In the study by Carter *et al*. [27], logistic regression, a multi-layer perceptron (neural network), and eXtreme gradient-boosted decision tree (XGBoost) models were trained to predict if missense mutations in *pncA* conferred resistance to pyrazinamide. The train/test dataset comprised 664 non-redundant missense amino acid mutations in the *pncA* gene, each of which had a label of either resistant or susceptible derived from the associated DST data. The dataset was built by combining mutations from an *in vitro* / *in vivo* mutagenesis study by Yadon *et al*. [13] with mutations from the first edition of the WHO catalogue of resistance-associated mutations [33]. The validation dataset was built by aggregating previously-published clinical samples which had both a missense mutation in *pncA* and a DST result, however the reported performance on this dataset was poor [27], in part due to the variability in the DST results.

While these models included some structural features, traditional machine learning models, due to the constraints of having a tabular input, cannot easily access information embedded in the full 3-dimensional structure of the protein. Another limitation was that only PncA sequences with a single missense mutation could be used; any samples with two or more mutations were discarded. As a result, the models are limited to predictions based on the presence of only a single mutation in the resistance-associated gene *pncA*.

Deep learning has been successfully applied to many problems in computational biology [34]. Geometric deep learning methods, like graph neural networks (GNNs), have the significant benefit of being able to handle protein structural data as a 3-dimensional representation [35]. Graph convolutional networks (GCNs) [36] allow convolution operations, originally developed for image data processing, to be applied to molecular graph representations of proteins [37]. Such models can therefore extract interaction-aware features from nodes and their neighbours in a network. This can be useful in protein representation learning since a degree of contextual information can be captured from an amino acid’s neighbouring residues in 3-dimensional space, as these relationships are expressed in the network’s connectivity. Adding further information, for example edge weights based on the distance between two residues in the protein structure, can further enrich the graph representation of the protein. Graph neural networks naturally account for the translational and rotational symmetries of molecular structures and stacking multiple GCN layers allows for information to be propagated through the graph so that relationships between distant residues can be learned.

Whilst GCNs show significant potential as a method for learning which mutations confer resistance to an antibiotic, they require sufficient genetic variability and a large dataset for training, due to their inherent expressivity. The slow mutation rate of 0.2-0.5 single nucleotide polymorphisms (SNPs) per year of *M. tuberculosis* [38] ensures that the overall genetic variability is low; this made it easier to apply tabular machine learning methods [12, 27]. In addition, the current gap in resistance prediction for TB can be small for some drugs, as catalogues generally perform well, especially for drugs with well-defined “resistance-determining regions” [39]. For pyrazinamide, an added problem is the inherent unreliability of the DST methods used to determine the resistance phenotype of *M. tuberculosis* isolates [40, 41]. This means the labelling error is high in training data which places an upper limit on the performance of machine learning models. We can therefore expect the performance improvements from using GCNs for resistance prediction in TB to be smaller than in other pathogens which don’t suffer from these shortcomings.

In this paper, we show that a GCN can accurately predict pyrazinamide resistance in PncA and has comparable performance to the best performing tabular machine learning methods when using the same dataset and train/test split as Carter *et al*. We then adopt two more train/test split strategies; split by amino acid position, and by structural cluster, to investigate how well the GCN and the baseline methods in Carter *et al*. can generalise and predict resistance of mutations in unseen parts of the protein. We show that for these more difficult classification tasks, a GCN either outperforms or matches the methods in Carter *et al*., and largely maintains performance relative to the original train/test split. Whilst some studies have applied GCNs to AMR prediction [42–44], we believe this is the first to do this by explicitly modelling the structure of the protein in the graph. We hope this study serves as a proof of concept for applying GCNs to other pathogens, for example *E. coli* [45], where a greater degree of genetic variation constrains current prediction methods, and therefore the potential benefit of using expressive methods like GCNs, are comparatively greater.

## Methods

A full end-to-end overview of the methods used to train the GCNs presented in this paper can be seen in Fig. 2.

**Fig. 2:**
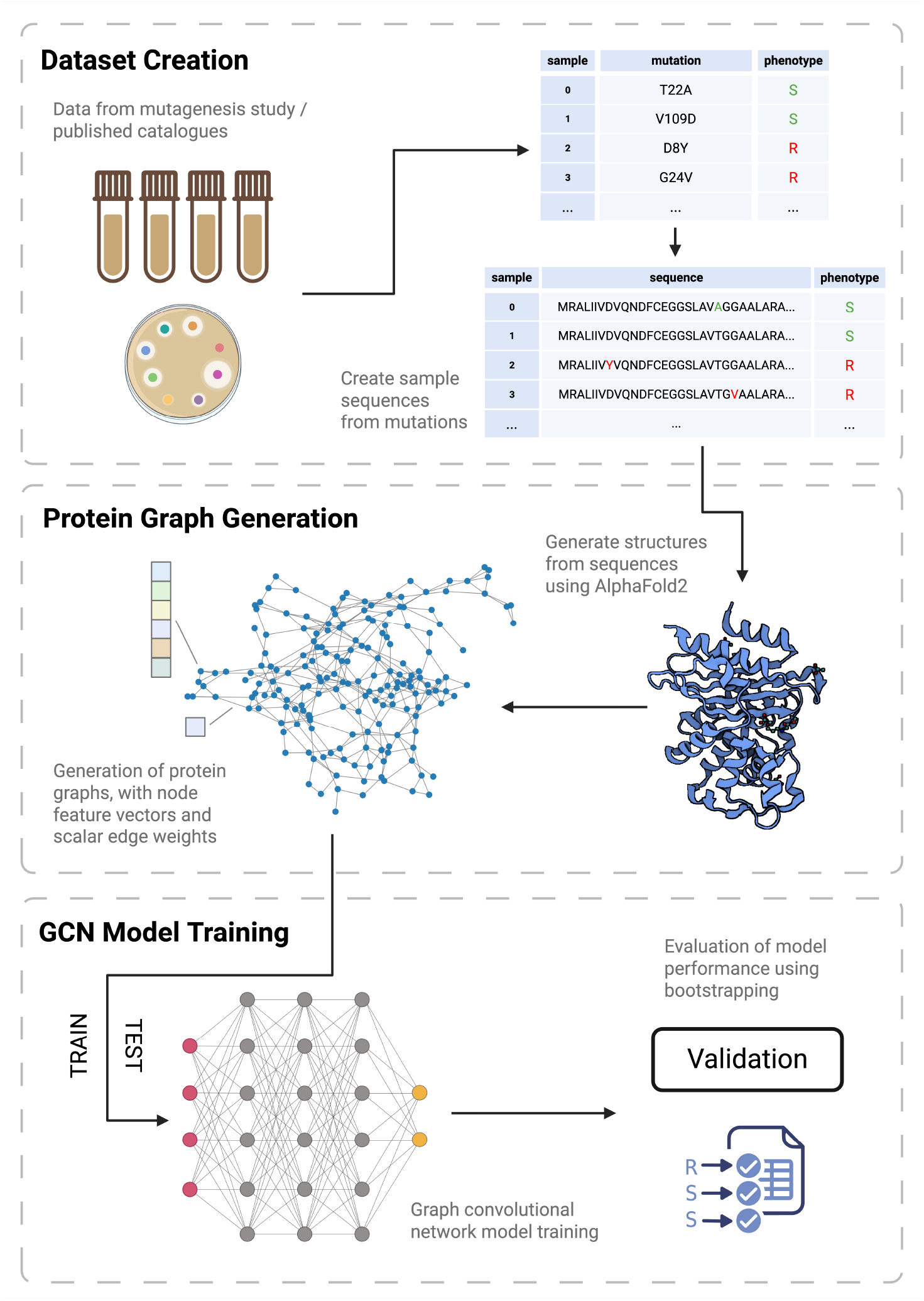
Overview of the workflow from dataset creation to model training and validation. The test/train dataset from a previous study was used to facilitate comparison [27]. Mutations from published catalogues and a mutagenesis study are used to create a dataset of PncA sequences with phenotype labels. Predicted structures of these sequences are then generated using AlphaFold2, and protein graphs are constructed to be used as inputs to the GCN. Each node is attached with a node feature vector and each edge is attached with an edge weight. The GCN model is trained on the train dataset and then evaluated using bootstrapping (*n=10*) on the test set.

**Fig. 3:**
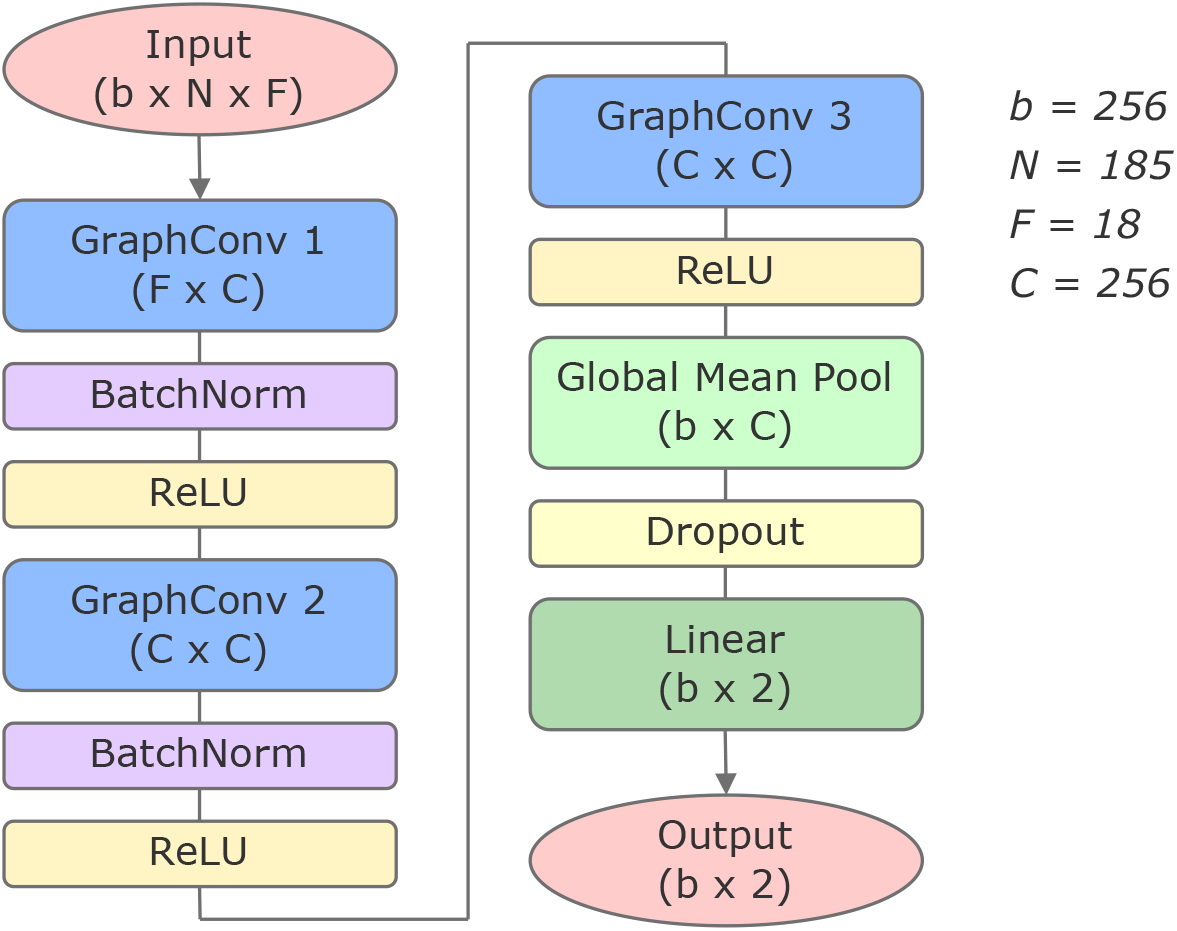
Architecture of layers in the GCN model. Dimension sizes are shown for the GCN trained using the Carter *et al*. [27] dataset split. The batch size is *b, N* represents the number of nodes in the graph whilst *F* is the number of node features. *N* is defined by the number of residues in the sequence, and as such is constant for all samples in the dataset (as they each contain a single amino acid substitution). The node features that comprise *F* can be seen in Table S1. The output dimension size for that layer, or the number of hidden channels for a given layer, is denoted by *C. b* and *C* were selected for by hyper-parameter tuning. Details of the parameter ranges used can be seen in Table S2.

### Dataset

The train and test datasets were downloaded from the GitHub repository [46] that accompanied the study by Carter *et al*. and consisted of 664 unique missense mutations randomly split 70:30 (Table 1). From the list of mutations, we constructed an equivalent dataset of PncA alleles, each containing a single mutation incorporated into the wild-type PncA sequence.

**Table 1:** Sample counts across dataset splits and phenotypes (labels). The train set and test set counts are shown for the original dataset split from Carter *et al*. [27], the *Amino Acid Position Split*, and the *Structural Cluster Split*. The ratio of susceptible (S) and resistant (R) samples is approximately maintained in the train and test set for each dataset split.

|  |  | Dataset Split |  |  |  |  |  |
| --- | --- | --- | --- | --- | --- | --- | --- |
|  |  | Carter *et al.* |  | Amino Acid Position |  | Structural Cluster |  |
| Phenotype | All | Train | Test | Train | Test | Train | Test |
| All | 664 | 464 | 200 | 462 | 202 | 463 | 201 |
| S | 315 | 218 | 97 | 227 | 88 | 217 | 98 |
| R | 349 | 246 | 103 | 235 | 114 | 246 | 103 |

We used this dataset of 664 *pncA* sequences to generate predicted structures using the ColabFold [47] implementation of AlphaFold2 [48]. For each input sequence from our constructed dataset, AlphaFold generated five structures ranked by the average predicted Local Distance Difference Test (pLDDT) - an estimate of the confidence in position of each residue. For each allele, we selected the top-ranked structure.

Since the *M. tuberculosis* structure of PncA [49] does not contain the structure of bound pyrazinamide we followed Carter *et al*. [27] and structurally fitted each of the AlphaFold2 predicted structures onto the experimentally-determined structure of *Acinetobacter baumannii* PncA [50], retaining the coordinates of the bound pyrazinamide and the Fe^2+^ ion in each case (Algorithm 1 in Supplement).

We created two more dataset splits for further experiments (Table 1). The *Amino Acid Position Split* was created by assigning samples to the train or test set based on the position of the residue in which their respective mutation is located. The *Structural Cluster Split* was created by using K-means clustering on the coordinates of the centres of mass of the residues in the wild-type PncA, then assigning samples to their respective cluster in a similar way to the *Amino Acid Position Split*. Assignment to the train or test set was then performed by the assignment algorithm (Algorithm 2 in Supplement). These methods are discussed further in the Results section.

### Protein Graph Representation

Protein graphs were derived from AlphaFold2’s predicted structural coordinates. Nodes in each graph corresponded to amino acid residues with edges indicating that the two connected residues lay within a defined distance, *d* (4 ≤ *d* ≤ 18 Å), that was explored during hyper-parameter tuning (Table S2). A distance of 12 Å between nodes was found to be the optimal threshold for defining an edge, where a given node’s position was specified by the centre of mass of the atoms of that residue. Self-loops were not included as edges.

Each node was annotated with features derived using sbmlcore [51], a Python package that calculates amino acid features for structural machine learning tasks. Features were chosen to best encode an amino acid’s chemical and structural properties and/or the effect that the amino acid substitution has on the protein’s function, in cases where that residue has been mutated. Along with features based on the amino acid’s chemical properties, distances of each amino acid to the pyrazinamide molecule and Fe^2+^ ion were included (Table S1). Nodes corresponding to mutated residues had scores from meta-predictors (DeepDDG [52], RaSP [53], MAPP [54] and SNAP2 [55]) attached. All other (wild-type) nodes were given a value representing a neutral or no change in the prediction of these features (-100 for SNAP2 and 0 for DeepDDG, RaSP and MAPP). Node features were normalised using `MinMaxScaler`. The scaler was fit only on the train set (to avoid data leakage) and then applied to the full train and test set.

An edge weight was attached to every edge in the graph. Edge weights were derived from distances between the centre of masses of residues in the PncA structure. Determining the edge weight calculation method from a choice of options (Table 2) was done as part of the hyper-parameter tuning sweep. Exponentially decaying edge weights were found to perform best. The value of *λ* was also optimised during hyper-parameter tuning.

**Table 2:** Edge weight calculation methods. The edge weight between nodes *i* and *j* is *w*(*i, j*), **r**_*i*_, **r**_*j*_ ∈ ℝ^3^ represent the 3-dimensional position vectors of the centres of mass of residues *i* and *j* and *cutoff* is the predefined cutoff distance such that if ∥**r**_*i*_ − **r**_*j*_∥ *< cutoff*, then an edge exists between **r**_*i*_ and **r**_*j*_. *λ* is a positive constant.

| Method | Calculation ($w(i, j)$) |
| --- | --- |
| None | 1 |
| Distance | $\ \mathbf{r}_i - \mathbf{r}_j\ $ |
| Inverse Distance | $\frac{1}{\ \mathbf{r}_i - \mathbf{r}_j\ }$ |
| Linear Decay | $1 - \frac{\ \mathbf{r}_i - \mathbf{r}_j\ }{\text{cutoff}}$ |
| Exponential Decay | $e^{-\lambda \cdot \ \mathbf{r}_i - \mathbf{r}_j\ }$ |

### Graph Convolutional Network

GCNs were implemented using the PyTorch Geometric library [56] with the protein graphs as inputs. We use the graph convolution layers `GraphConv`, based on Morris *et al*. [57]. Each layer takes as an input the adjacency matrix **A** and the node-level embeddings from the previous layer. The `GraphConv` operation aggregates node embeddings from a node’s neighbours and combines it with its own embedding to produce updated node embeddings for the next layer. The graph convolution operation for the input feature vector or embedding of a given node **x**_*i*_, and the resulting embedding 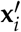, is defined as

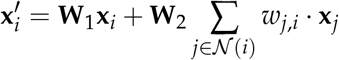

where **W**_1_ and **W**_2_ are learnable weight matrices, 𝒩 (*i*) is the set of all node neighbours of *i*, and *w*_*j,i*_ is the edge weight between two given nodes *j* and *i*.

The main GCN architecture consisted of three convolution layers followed by a fully connected linear layer for final classification. The use of three convolution layers each followed by ReLU is standard practice for GCNs designed for protein structure and similar applications [37]. `BatchNorm` layers were incorporated after each of the first two convolutions to improve stability and smoothness of training. The model was trained using the `AdamW` optimiser and `CrossEntropyLoss`, to output a binary classification of resistant or susceptible. Model tuning was performed using Weights & Biases [58] which efficiently runs hyper-parameter tuning sweeps and tracks experiments. The number of hidden layers, dropout probability, learning rate and weight decay were all selected by iteratively performing random sweeps. In all tuning sweeps, we maximised the F1-score of the test set.

We compared the GCN performance and that of several simple machine learning models (logistic regression, a simple neural network and gradient-boosted decision tree) as reported by Carter *et al*. [27]. The bootstrapping method from that study was replicated using the same random seeds to ensure that the same samples were included in each bootstrap.

## Results

### A GCN model can accurately predict pyrazinamide resistance

The GCN was trained on the train dataset and hyper-parameter tuning sweeps were iteratively run using a random search strategy. The model was evaluated using bootstrapping (*n=10*) on the test set. This model was compared to the previously published results of three tabular machine learning methods that were trained (and tested) on the same datasets: logistic regression (LR), XGBoost (XB), and a single layer neural network (NN) [27]. Of these, the XB model performed best on the test set (Table 3) and after evaluating with bootstrapping (Fig. 4).

**Table 3:** The performance of the GCN compared to three published machine learning models from Carter *et al*. [27]. All models used the same dataset and train/test split used in Carter *et al*. The results from the GCN model we present in our study is marked by (^†^) in the *Model* column.

| Model | Test F1 (%) | Test Sensitivity (%) | Test Specificity (%) | Test ROC AUC (%) | TN | FP | FN | TP |
| --- | --- | --- | --- | --- | --- | --- | --- | --- |
| LR | 77.4 | 79.6 | 72.2 | 82.8 | 70 | 27 | 21 | 82 |
| NN | 70.6 | 75.7 | 58.8 | 75.1 | 57 | 40 | 25 | 78 |
| XB | 80.2 | 80.6 | 78.4 | 82.2 | 76 | 21 | 20 | 83 |
| GCN<sup>†</sup> | **81.6** | **81.6** | **80.4** | **82.8** | **78** | **19** | **19** | **84** |

**Fig. 4:**
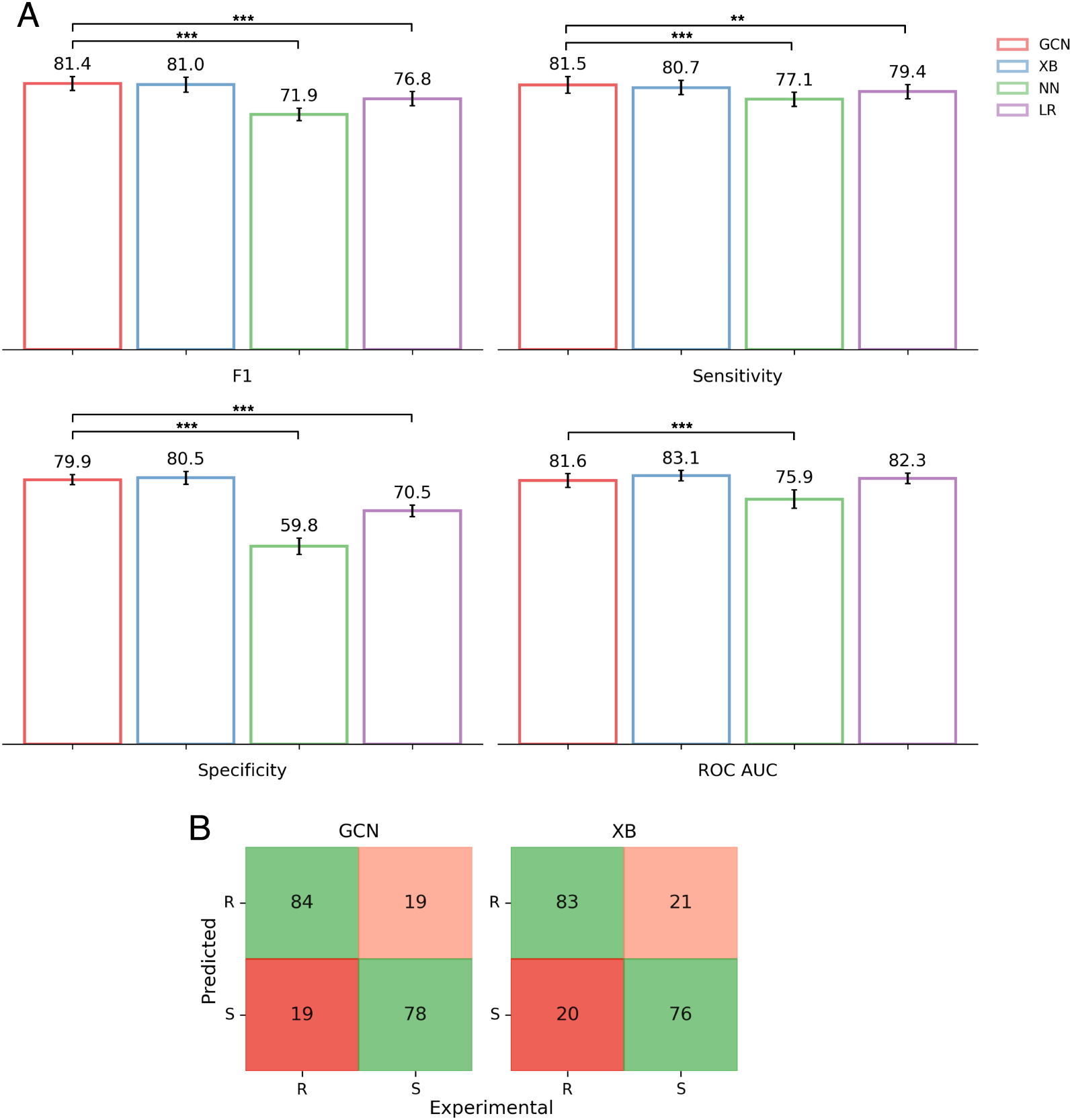
GCN outperforms or matches the performance of the classical tabular machine learning methods from Carter *et al*. [27]. **(A)** The GCN model performance on the bootstrapped test set *(n=10)*. The performance of XB, NN and LR taken as reported in Carter *et al*. Error bars represent 95% confidence intervals, reflecting the distribution of scores across bootstrap resamples. Statistical significance is assessed (paired *t*-test) on matched bootstrap resamples. Brackets represent significant differences relative to the GCN model (^∗^ : *p ≤* 0.05; ^∗∗^ : *p* ≤ 0.01; ^∗∗∗^ : *p ≤* 0.001). **(B)** Confusion matrices of the GCN model for the test set, compared to XB. Very Major Errors (VME) are coloured red and are considered worse than Major Errors (ME), which are coloured orange.

Our GCN model achieved higher scores in all performance metrics than XB on the test set with a sensitivity of 81.6 % and a specificity of 80.4% (Table 3). The GCN performed equivalently well to XB after bootstrapping (Fig. 4A, paired *t*-test, all metrics *p* ≥ 0.05, hence no significant difference) and significantly better than both the NN and LR models for all four performance metrics (Fig. 4A) except ROC-AUC, where it was only significantly better compared to the NN model. The GCN had two fewer major errors (ME) and one fewer very major error (VME) in the test set compared to the existing XB model (Fig. 4B). MEs are defined as susceptible samples incorrectly classified as resistant (false positives) and VMEs are resistant samples incorrectly classified as susceptible (false negatives) [59]. The distinction is made as there is typically a worse clinical outcome if a resistant sample is incorrectly classified rather than *vice versa*, as MEs result in overtreatment, e.g. a stronger drug than is needed being used as treatment, whilst VMEs could result in an ineffective drug being used, treatment failure and spread of the resistant strain.

### GCN successfully generalises to predict resistance of samples with mutations in unseen positions and structural regions

A random split was used by Carter *et al*. [27], with a fixed random seed for reproducibility, which we initially used to train the GCN and compare performance to the methods published in that study. However, this resulted in samples with mutations at the majority of amino acid positions ending up in both the train and test sets (Fig. 1B). This could mean data leakage and the train/test split containing unintended biases [60, 61], potentially causing the model to memorise trends at given amino acid positions in the train set.

We further investigated the generalisability of the GCN’s ability to predict pyrazinamide resistance by considering alternative methods to split the dataset (Table 1). We did this by ensuring that the train and test sets each contained samples with mutations at distinct positions (Fig. S5). This meant that the test set exclusively featured mutations at positions that the model had never seen during training. Assignment of amino acid positions to the train and test sets was performed by a greedy algorithm with random initialisation (Algorithm 2 in Supplement). We refer to this split as the *Amino Acid Position Split*. Test set performance therefore reflected the model’s ability to generalise and predict resistance based on mutations at unseen positions.

We tested this further by using an additional dataset split that provided another more difficult learning task. By dividing PncA into regions based on its 3-dimensional structure, we use these regions to create the train/test split and ensure that each set contained samples with mutations from structurally distinct regions (Fig. 5). We refer to this as the *Structural Cluster Split*. To create this, we took the coordinates of the centres of mass of the residues in the wild-type PncA structure and used K-means clustering to divide residues into their respective clusters (Fig. 5A). Since all samples in the dataset contain a single amino acid substitution, we can assign each sample to the cluster that contains the residue at which their mutation is found. Whilst K-means clustering is agnostic of any biological significance, such as secondary structure or protein function, it provides a simple, unbiased method for splitting the protein into different structural regions. We chose a value of *k=18*, as this gave appropriately sized clusters of residues. Assignment of structural clusters to the train and test sets (Fig. 5B) were performed by the same algorithm used to create the Amino Acid Position Split (Algorithm 2 in Supplement). In this scenario, the model now has to generalise to unseen structural regions of the protein, not just unseen positions (Fig. S5), and tests the GCN’s capacity to learn deep representations of node neighbourhoods.

**Fig. 5:**
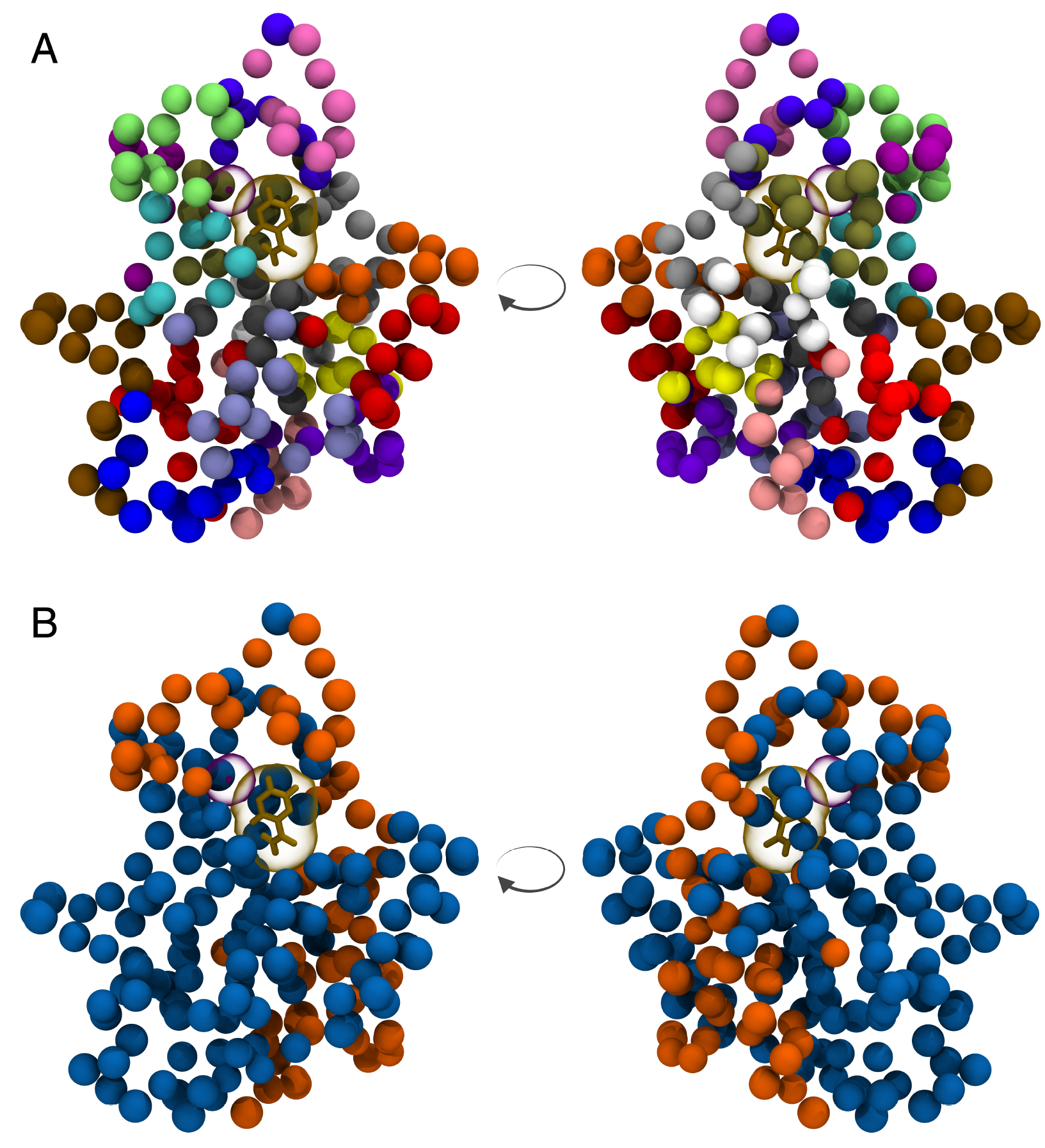
Structural clustering of PncA by K-means and assignment of samples to the train and test sets in the *Structural Cluster Split*. PncA structure represented with each residue’s C-*α* atom shown as a bead, with bound pyrazinamide and the Fe^2+^ shown (as in Fig. 1). Structure rendered using VMD [26]. The second image in each panel shows the protein rotated 180°. **(A)** K-means clustering assigns each residue in PncA to a cluster (*k=18*). Each cluster is represented by a different colour. Samples in the dataset are assigned to the cluster that contains the position of their mutated residue. **(B)** Each cluster is assigned to either the train or test set using the assignment algorithm (Algorithm 2 in Supplement). Samples are assigned to the train set if they have a mutation at a residue coloured blue, and to the test set if they have a mutation at a residue coloured orange.

We trained and evaluated GCNs on these two new dataset splits using the same hyper-parameter tuning and bootstrapping method as described previously. We also used code from the GitHub repository [46] that accompanied the work from Carter *et al*. to run the LR, XB and NN models on these new dataset splits.

Our results now show that, with these more difficult learning tasks, the GCN achieves a significantly higher F1 score (paired *t*-test) than the traditional methods for both dataset splits (Table 4). For the *Amino Acid Position Split*, the GCN achieved a mean bootstrapped test F1 score of 83.9 *±* 2.1%, significantly outperforming LR (81.9 *±* 1.6%), XB (77.5 *±* 2.2%) and NN (73.2 *±* 1.6%). In addition, the GCN demonstrated balanced performance across sensitivity (81.4 *±* 3.1%) and specificity (83.2 *±* 2.4%), and achieved the highest ROC AUC (86.2 *±* 1.5%) among all models for this split.

**Table 4:** GCN demonstrates more reliable generalisability than the selected traditional machine learning methods. GCN outperforms machine learning methods from Carter *et al*. [27] when the dataset is split using either the amino acid position based or the structural cluster based method. The same samples were used in each bootstrap for evaluation of all models. Code from Carter *et al*. GitHub [46] was used to run the LR, XB and NN baselines on the two new dataset splits. The results from the GCN model we present in our study is marked by (^†^) in the *Model* column. Values are reported as mean *±*95% confidence intervals, reflecting the distribution of scores across bootstrap resamples. Statistical significance is assessed (paired *t*-test) on matched bootstrap resamples. Asterisks represent significant differences relative to the GCN model (^∗^ : *p* ≤ 0.05; ^∗∗^ : *p* ≤ 0.01; ^∗∗∗^ : *p* ≤ 0.001).

| Dataset | Model | Bootstrapped<br>Test F1<br>(%) | Bootstrapped<br>Test<br>Sensitivity<br>(%) | Bootstrapped<br>Test<br>Specificity<br>(%) | Bootstrapped<br>Test<br>ROC AUC<br>(%) |
| --- | --- | --- | --- | --- | --- |
| Amino<br>Acid<br>Position<br>Split | LR | $81.9 \pm 1.6^{**}$ | $82.5 \pm 2.8$ | $74.9 \pm 3.1^{***}$ | $85.0 \pm 1.8$ |
| | NN | $73.2 \pm 1.6^{***}$ | $70.7 \pm 2.5^{***}$ | $70.1 \pm 3.4^{***}$ | $79.3 \pm 1.4^{***}$ |
| | XB | $77.5 \pm 2.2^{***}$ | $73.6 \pm 3.0^{***}$ | $78.3 \pm 3.2^{**}$ | $81.4 \pm 1.9^{***}$ |
| | GCN<sup>†</sup> | **$83.9 \pm 2.1$** | $81.4 \pm 3.1$ | **$83.2 \pm 2.4$** | **$86.2 \pm 1.5$** |
| Structural<br>Cluster<br>Split | LR | $77.0 \pm 1.6^*$ | $71.5 \pm 2.1^{***}$ | $85.8 \pm 1.2^{***}$ | $86.0 \pm 1.2$ |
| | NN | $65.5 \pm 1.8^{***}$ | $62.5 \pm 2.0^{***}$ | $71.3 \pm 3.6^{***}$ | $74.5 \pm 1.9^{***}$ |
| | XB | $73.3 \pm 2.3^{***}$ | $62.9 \pm 2.8^{***}$ | **$91.3 \pm 1.3^{***}$** | **$86.2 \pm 1.5$** |
| | GCN<sup>†</sup> | **$79.4 \pm 1.9$** | **$78.4 \pm 2.7$** | $80.7 \pm 1.7$ | $85.7 \pm 1.1$ |

For the more challenging *Structural Cluster Split*, the GCN again achieved the highest F1 score at 79.4 *±* 1.9%, significantly outperforming LR (77.0 *±* 1.6%), XB (73.3 *±* 2.3%) and NN (65.5 *±* 1.8%). While XB achieved a higher specificity (91.3 *±* 1.3%), this came at the expense of substantially lower sensitivity (62.9 *±* 2.8%), whereas the GCN maintained a more balanced trade-off between sensitivity (78.4 *±* 2.7%) and specificity (80.7 *±* 1.7%). The GCN also achieved a ROC AUC of 85.7 *±* 1.1%, matching the best-performing baseline models (Table 4).

### Node features and graph structure both contribute to GCN performance

Next we investigated the relative effects of node features and graph structure in our GCN model by performing a series of ablation experiments. The meta-predictor features (SNAP2, DeepDDG, MAPP and RaSP) were included as node features as they are expressive measures from other neural network models that aim to quantify the impact a mutation has on a protein. SNAP2 estimates the effect on function due to a mutation [55] whilst DeepDDG and RaSP predict how a point mutation in a protein affects its stability [52, 53], and MAPP quantifies the evolutionary constraints imposed on a given position in a protein [54]. It is therefore likely that, as node features, these could provide important information about whether a certain mutation could result in a resistant or susceptible phenotype. These features were also found to be four of the top five most discriminatory individual features in the previous study [27], as measured by the ROC AUC of univariable logistic regression models trained on each of the individual features. However, these meta-predictor features could have been influencing the GCN model by, in effect, flagging where in the protein a mutation has occurred, as the meta-predictors’ values were only attached to mutated residues (with wild-type residues all having the same zero-equivalent value).

To investigate the relative contribution of these features to the GCN, using the Carter *et al*. [27] dataset split, we trained and tested a GCN using just SNAP2, DeepDDG, MAPP and RaSP (meta-predictor features - MPFs) as node features (Fig. 6 - “MPFs only”). We also trained a model having excluded these four features, hence the model only had available node features based on chemical properties and distances, (Fig. 6 - “No MPFs”) and compared performance to the full GCN model. Both models performed significantly worse than the full GCN.

**Fig. 6:**
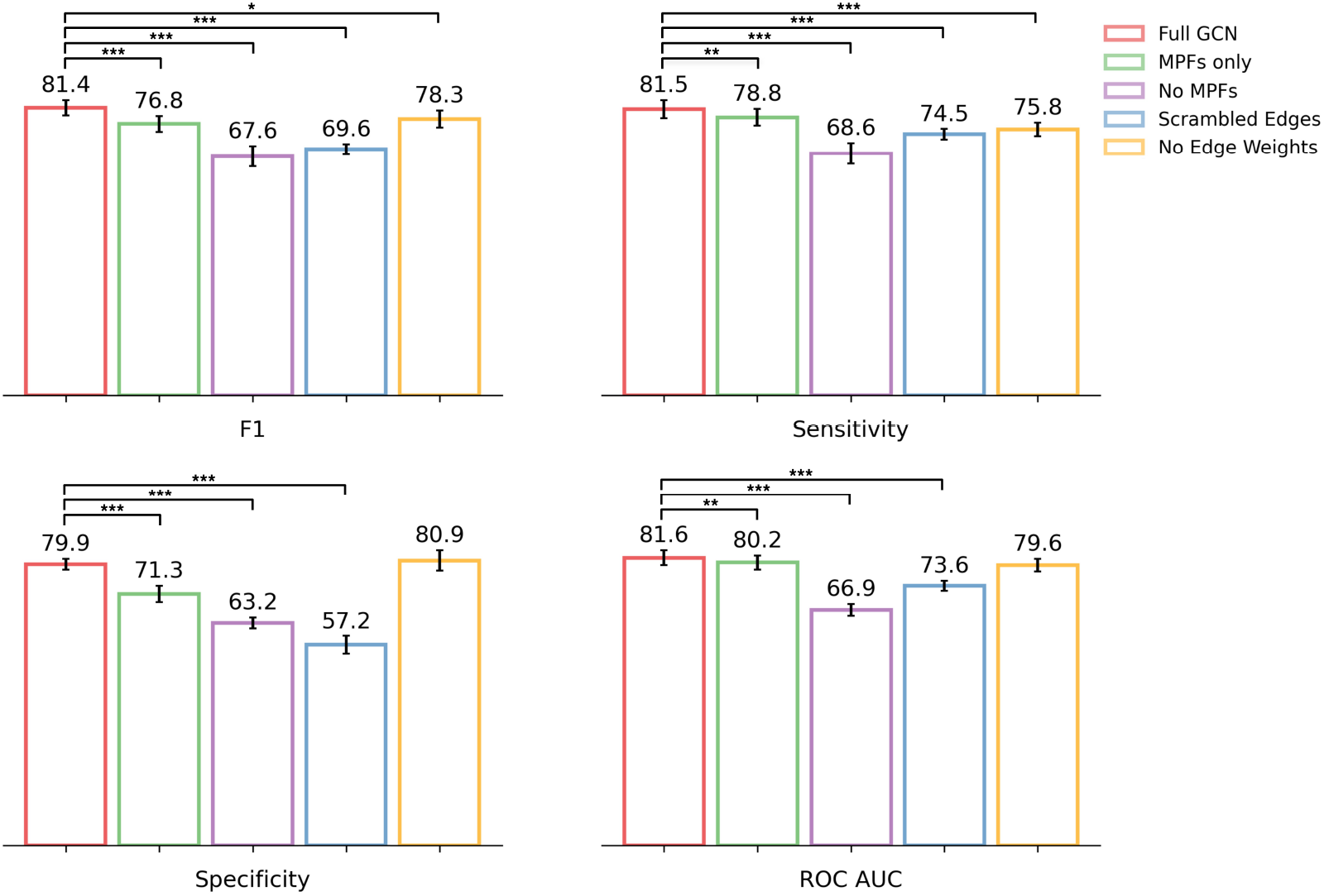
Ablation of different aspects of node features or graph structure results in a significant reduction in GCN performance. GCN control experiment models performance on bootstrapped test set *(n=10)*. Error bars represent 95% confidence intervals, reflecting the distribution of scores across bootstrap resamples. Statistical significance is assessed (paired *t*-test) on matched bootstrap resamples. Brackets represent significant differences relative to the GCN model (^∗^ : *p* ≤ 0.05; ^∗∗^ : *p* ≤ 0.01; ^∗∗∗^ : *p* ≤ 0.001).

A major advantage provided by graph machine learning is the capacity to represent the protein structure through the connectivity of the graph. We evaluated the contribution of the graph structure to model performance by training a GCN with protein graphs containing scrambled edges (and a full set of node features). Edge scrambling was implemented by maintaining the source node of each edge and then randomly shuffling the target node and edge weight. This avoided totally destroying the graph network by keeping the the global number of edges and node degree distribution unchanged. Any reduction in performance therefore reflected a loss of meaningful topological structure. The GCN model trained on scrambled graph (Fig. 6 - “Scrambled Edges”) exhibited lower performance, from which we can infer that the spatial and structural information being represented by the graph contributes positively to performance.

For each ablation experiment, we performed independent hyper-parameter sweeps, although the optimal parameters remained consistent across setups. This suggests that the drop in performance was caused by the omission of key features or the disruption of graph connectivity, not a lack of optimisation in hyper-parameter space.

### Meta-predictors, particularly DeepDDG, were the features that most influenced the GCN’s predictions

To gain additional insight into the relative effect that each node feature has on the model inference we applied a feature attribution method. This calculates importance scores for a given feature, representing the contribution of that feature to the model’s prediction output [62]. Gradients of the model prediction with respect to the input features reflect the change in output in response to small changes in the input, and hence can be a good approximation for explainability of a model’s feature importance, akin to saliency maps in image classification [63]. For the GCN model trained on the Carter *et al*. dataset split [27], we implemented a gradient-based feature attribution method which calculated the gradients of the output logits for the predicted class with respect to the input features, for inferences made on samples in the test set. The DeepDDG score had the highest relative feature importance followed by the scores of RaSP, MAPP and Snap2 (Fig. 7). DeepDDG predicts the change in stability of a protein, measured by the change in Gibbs free energy (ΔΔ*G*), as a consequence of a missense mutation [52]. Protein stability is often connected to function, as a loss in stability is likely to result in a loss or alteration of enzymatic function. A reduction in the enzymatic activity of PncA would prevent activation of the pro-drug pyrazinamide and could therefore lead to resistance [14]. Therefore, it is reasonable to hypothesize that a large change in ΔΔ*G* of a protein results in a resistant phenotype. A molecular dynamics study also found that significant differences in ΔΔ*G* between wild-type and mutant PncA can be a useful predictor of pyrazinamide resistance [64].

**Fig. 7:**
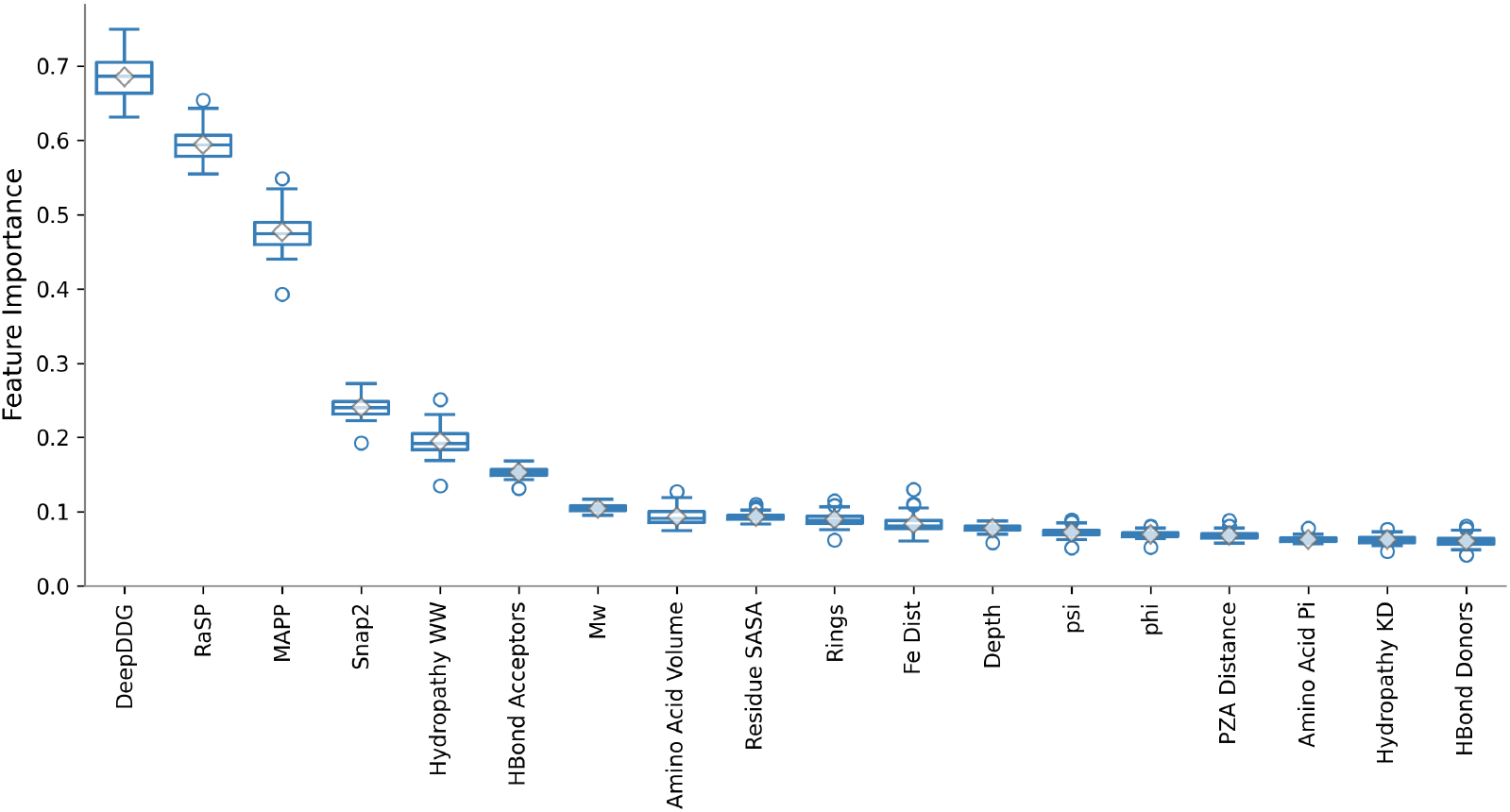
Gradient-based feature attribution showed meta-predictor features had the most influence on the model’s output. Box plots of feature importances of each feature across the test set. Hydropathy WW/KD apply the Wimley–White and Kyte-Doolittle residue hydrophobicity scales, respectively, to each residue; SASA = solvent accessible surface area; Hbond Acceptors/Donors count the number of hydrogen bond acceptors and donors on each sidechain; Mw = molecular weight; Rings is a count of the number of aromatic rings in the sidechain; psi and phi are the backbone Ramachandran angles

Aside from meta-predictor features, whilst the remaining node features generated by `sbmlcore` individually contribute only a little to the model output, together they are predictive. This is shown by the comparative performance of GCNs trained with differing sets of features, which demonstrates that the features based on the amino acid’s chemical properties and structural distances are still predictive (Fig. 6). Distances (from either the bound pyrazinamide or catalytic Fe^2+^) did not have noticeably higher feature importances than other `sbmlcore` features (Fig. 7). This is perhaps surprising as one would expect the distance of a residue from the bound drug or the catalytic Fe^2+^ ion to inversely correlate with how likely a mutation at that position is to result in a resistant or susceptible phenotype. For example a perturbation is likely to have more impact on the binding of the drug if it occurs to a residue near the active site, or if it is close enough to directly disrupt the catalytic Fe^2+^ ion [65]. One possible reason as to why the distance features did not exhibit higher importances could be that this distance information is implicitly represented in the graph structure or even in the meta-predictors themselves.

The evaluation of individual feature performance by univariable logistic regression by Carter *et al*. also found Snap2 (81% AUC), DeepDDG (80% AUC), MAPP (74% AUC) and RaSP (73% AUC) to be the 1st, 2nd, 3rd and 5th best performing features respectively, where features were selected to be used in models if they achieved 55% AUC or greater [27].

### Edge weights aid the GCN when making an inference

Next we investigated the effect that edge weights have on GCN training and performance. The `GNNExplainer` class from PyTorch Geometric provides algorithms that can generate explanations to help understand why a GNN makes a certain decision. We used `GNNExplainer` to evaluate edge importances which are calculated by masking each edge in the graph in turn and measuring the effect this has when the model subsequently makes an inference. This was run on every sample in the test set and an average edge importance was calculated for each edge, as denoted by the combination of the source and target node indexes. The frequency of each edge occurring across the dataset is not considered (although we expect this to be very similar for all samples). The average edge importances of the GCN trained on the Carter *et al*. dataset split [27] showed a flat distribution (Fig. 8A). We hypothesised that this is because the edge weights were informing the model, with sufficient accuracy, about the relative value of each edge, so the edge masking algorithm of `GNNExplainer` suggested that the GCN was not employing any further bias when making an inference.

**Fig. 8:**
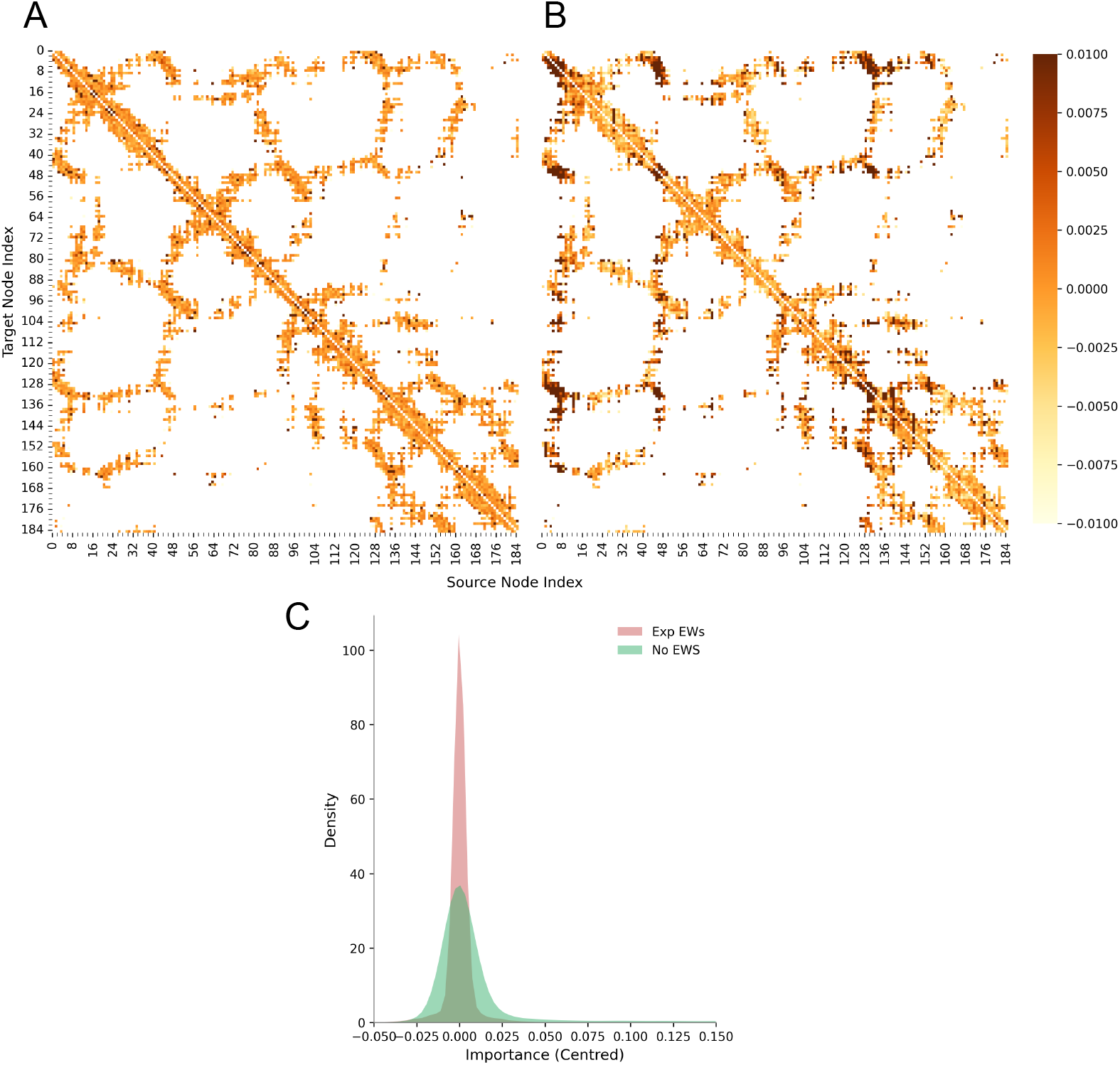
Greater variability in average edge importance is observed for the GCN model when trained without edge weights. Heatmaps show the average edge importance across all PncA graphs in the test set for each given edge. Average edge importances are centred around the median for each model respectively. Edge importance is obtained from the edge mask calculated by GNNExplainer for the GCN trained with edge weights **(A)** and without edge weights **(B). (C)** Kernel density estimate plot of the edge importances of edges in the GCN model with edge weights (red) and without edge weights (green). There is a significant difference between the two distributions (Wilcoxon signed-rank test; *p* ≤ 0.001).

As a comparison, we trained a GCN without edge weights and evaluated the edge importances. The edge cutoff distance (12 Å) was kept the same so that the corresponding protein graphs in the datasets of both models have identical edge indexes. The GCN without edge weights achieved significantly lower performance, notably an F1 score of 78.3 % (Fig. 6 - “No Edge Weights”), than the full GCN (with exponential decay edge weights, *λ* = 2, Table 2), whilst the average edge importances showed significantly more variability (Fig. 8C). This suggests that the absence of edge weights caused the GCN to bias some edges more than others when making an inference. The pattern in the heatmap of edge importances (Fig. 8B) can be seen in some regions to resemble corresponding regions in the heatmap of edge weights (Fig. S3) used in the full GCN. We hypothesise that the edge weights, which are based on inter-residue distances, are providing important information that the model is using, whilst the model trained without edge weights is, in some capacity, attempting to learn these relationships.

### AlphaFold2 predicted structures showed small deviations that were greater in resistant PncA variants than susceptible variants

Each protein graph is constructed based on the atomic coordinates of residues in the respective protein, as specified in the PDB file. In order to predict protein structures for every PncA variant, we used the AlphaFold2 implementation, ColabFold [47]. Given the importance of DeepDDG, we hypothesised that the structures of resistant alleles might be more distorted than those of susceptible alleles and hence calculated the structural variation between the predicted structures and the wild-type structure. To control for any bias introduced by AlphaFold2, we calculated the root mean square deviation (RMSD, over C*α* atoms) against an AlphaFold2 predicted structure of the wild-type sequence. For all 664 structures across both the train and test set, the mean RMSD was 0.107 Å (Table 5) and ranged from 0.031 Å to 0.540 Å, suggesting there was some predicted structural variability (Fig. S1).

**Table 5:** The root mean square deviation (mean *±* standard deviation, Å) across dataset splits and phenotypes of the C_*α*_ atoms in the AlphaFold2 predicted structures relative to an AlphaFold2 predicted wild-type structure.

| Phenotype | Dataset |  |  |
| --- | --- | --- | --- |
|  | Full | Train Set | Test Set |
| All | 0.107 ± 0.074 | 0.105 ± 0.069 | 0.111 ± 0.086 |
| S | 0.090 ± 0.069 | 0.089 ± 0.066 | 0.093 ± 0.075 |
| R | 0.122 ± 0.076 | 0.119 ± 0.069 | 0.127 ± 0.093 |

Interestingly the predicted structures of resistant alleles were more likely to have higher RMSD values than susceptible alleles (Fig. 9). We also note that this trend is largely observed across the whole sequence (Fig. S4). This suggests that mutations that make PncA resistant to pyrazinamide are more likely to lead to, on average, a greater perturbation to the protein’s structure. Such perturbations could directly affect the binding site, resulting in the disruption of pyrazinamide binding, or could affect the stability of the protein as a whole.

**Fig. 9:**
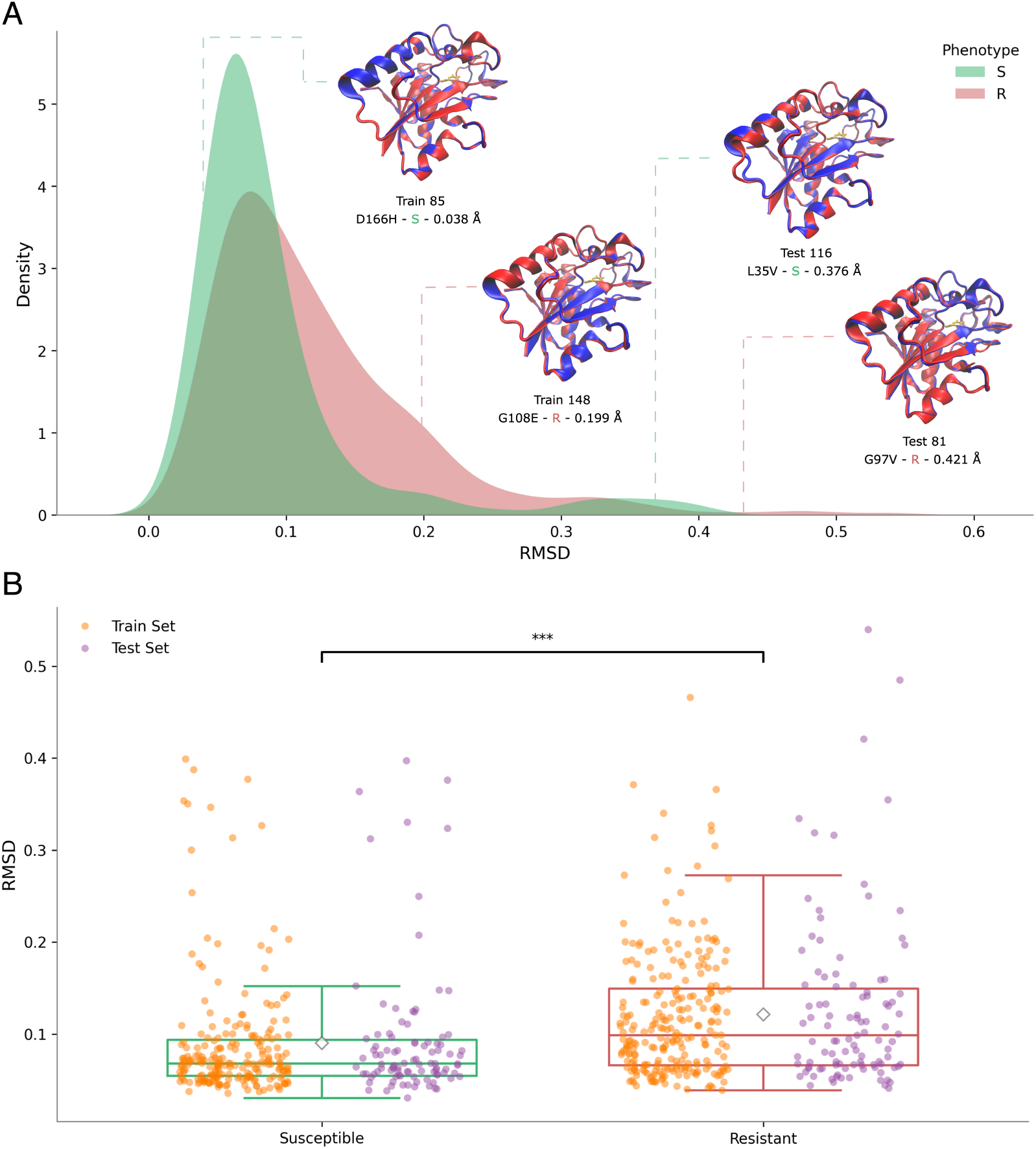
The AlphaFold2 predicted structures of resistant PncA alleles are more different to the wild-type sequence than susceptible alleles. **(A)** Kernel density estimate plot of the root mean square deviations (RMSD, Å) of predicted structures of susceptible (green) and resistant (red) PncA alleles. RMSD is measured over the C_*α*_ atoms with respect to the AlphaFold2-predicted wild-type structure. Selected structures of PncA variants (red) aligned with wild-type PncA (blue) are shown on the plot. **(B)** Box plot of RMSDs of susceptible (green) and resistant (red) PncA alleles, further split into whether they belong to the training (orange) or test (purple) sets. Resistant alleles have a significantly different RMSD distribution than susceptible variants (paired *t*-test; ^∗∗∗^ : *p* ≤ 0.001).

## Discussion

We have shown that a GCN, trained to predict whether PncA variants confer resistance to pyrazinamide, achieves comparable performance to the best performing tabular machine-learning models [27]. This is perhaps surprising as, whilst GCNs are powerful and parameter-heavy, they can be prone to overfitting and learning noise when trained on small datasets with little variation. In contrast, methods like XGBoost are well-suited for small to medium sized datasets, as we have here, and have a lower risk of noise amplification. GCNs are also more computationally expensive to train than XGBoost and other classical machine learning models and require slightly more time to make an inference. Inference time increases further when the generation of a structural prediction using AlphaFold2 (or an equivalent method) is included and accounted for. However, the GCN models we have trained are relatively small (Table S3), and therefore can be trained and run on modern hardware very easily without incurring significant computational expense.

We have demonstrated that the GCN is able to generalise and predict resistance in samples with mutations in unseen positions and structural regions. One interesting finding was that the GCN performed better with the *Amino Acid Position Split* than the random split from Carter *et al*. [27]. One reason for this could be that the random split was causing an unintended bias where, because the dataset was small, an imbalance arose in the phenotype labels for certain amino acid positions between the train and test sets. By splitting the data based on amino acid position of the mutation, during training the model could access either all of the labels for a given position or none of them, hence removing the risk of it learning a skewed distribution.

Whilst using K-means clustering to create the *Structural Cluster Split* has some shortcomings, notably the lack of biological significance of the clusters, the minimal drop in performance of the GCN demonstrates strong potential for this method to generalise successfully. Choosing an appropriate value of *k* was also important. A *k* that was too large meant the clusters would be too small and not represent a large enough region in the protein structure, whilst a *k* that was too small would result in very large clusters, a dataset split that was too aggressive and a learning problem that was too difficult.

There are other shortcomings in our approach. Better methods could be employed for attaching the drug to the AlphaFold2-predicted structures than by using a structural fit, as this assumes the mutations result in little to no perturbation to the binding site. Recent advancements in AlphaFold3, which can now perform structural predictions with ligand docking, will likely better represent the interaction between the drug and the protein [66]. This could also open the door to including the bound drug as a node(s), which might further refine our graph representation. That said, using AlphaFold2-predicted structures for each PncA allele may already provide most of the benefit by introducing variability in the graph connectivity as well as allowing for more accurate distance-based node features (distance to pyrazinamide or Fe^2+^) and protein secondary structure-based features. As discussed in the Results, a key limitation of our feature engineering is how we have attached meta-predictor features. Since the output of these models is only attached to the nodes representing the mutated amino acid, this could act as a flag to the model indicating where the site of the mutation is. This could bias the model to learn just the amino acid substitution and not infer from the protein as a whole when predicting the phenotype. Finally we could potentially improve our representation of amino acids in the GCN by using embeddings from protein language models (pLMs) such as ESM3 [67]. Future investigations into this should also involve benchmarking against using embeddings from pLMs on their own in classical machine learning methods.

One can expect that, with a larger dataset, GCNs will scale and generalise even better. In particular, we see significant potential benefit in the application of GCNs in predicting drug resistance in pathogens which intrinsically have greater genetic variability (including in allele length) than *M. tuberculosis*, for example in *E. coli* [68]. For *M. tuberculosis* genes that have well-defined “resistance-determining regions”, such as *rpoB* and *katG*, rules-based and catalogue methods already perform well. Mutations outside of this region are unlikely to have any effect on the resulting phenotype. Whilst the performance gap in these cases is small (sensitivity and specificity can be *>*95% [69]) and so the potential gain is limited, we would still expect GCNs (and other machine learning methods) to be able to at least match this performance [70]. GCNs can also capture certain elements that traditional machine learning methods cannot: this includes, but is not limited to, incorporating alleles with insertions and deletions, as well as the presence of multiple mutations in a single allele – all cases that were not included in our dataset so as to maintain a direct comparison with previous studies. It is our hope and intention to extend this GCN-based approach to learn, and therefore predict *de novo*, antimicrobial resistance in other pathogens.

## Conclusions

We demonstrate that a GCN can predict pyrazinamide resistance in *M. tuberculosis* by learning structural and chemical features in the context of the predicted structure of each *pncA* allele. Our GCN model achieves equivalent performance to the best performing traditional tabular machine learning methods, in spite of being trained on a small dataset with low variability. We also show that a GCN can generalise as effectively as or better than these traditional tabular methods and predict resistance in samples with mutations in unseen regions of the protein. Despite being more complex, GCNs have the potential to learn and predict the antimicrobial resistance profile of genes with a far greater degree of genetic variability than is observed in *M. tuberculosis*, potentially leading to significant improvements in both our understanding of resistance mechanisms and drug susceptibility testing for a far wider range of pathogens.

## Declarations

### Ethics approval and consent to participate

This work used a published, open dataset as described in Methods and hence no ethics approval or consent to participate was required.

### Consent for publication

No individual person’s data are included and hence no consent for publication is required.

### Availability of data and materials

An accompanying GitHub repository is available [71]: this contains the train/test dataset and Python code to retrain the model as well as states of the final models and code to redraw the main figures in this manuscript.

### Authors’ contributions

DD and PWF conceived of the study. DD, VMB & DA carried out the data preparation, model building, training and analysis steps with JAM & PWF providing advice. DD, JAM and PWF wrote the manuscript.

