## Supplemental Information for "Predicting pyrazinamide resistance in *Mycobacterium tuberculosis* using a graph convolutional network"

**Algorithm 1: Structural alignment of PncA variant to wild-type for attaching pyrazinamide.** Alignment performed using VMD. Target structure pdbs obtained from AlphaFold structural predictions.

Load wild-type structure .pdb as **reference**;
Load target structure .pdb from as **mobile**;
Select C$\alpha$ atoms within 4 Å of pyrazinamide in **reference** as **bindingsite**;
Define alignment atoms (C, C$\alpha$, N) in **reference** and **mobile** with **bindingsite**;
Align **mobile** to **reference** with `measure fit`;
Add pyrazinamide coordinates to **mobile**, as specified in **reference**

**Algorithm 2: Greedy assignment of groups to train and test sets.** Assigns samples with mutations in residues at either a given position or in a given structural cluster to the train or test set. Used to create the *Amino Acid Position Split* and the *Structural Cluster Split*.

Define group labels for each sample as **ids**;
  (amino acid positions or structural cluster identifiers);
Set desired test fraction $f = 0.3$ and random seed $s$;
Initialise random number generator with seed $s$;
Count number of samples assigned to each unique group in **ids**;
Randomly shuffle the list of groups;
Compute total number of samples $N$ and target test size $N_{\text{test}} = f \cdot N$;
Initialise empty set of test groups and running sample count $r = 0$;
**for** *each group g in shuffled group list* **do**
  **if** $r \geq N_{test}$ **then**
    break;
  Assign group $g$ to test set;
  $r = r +$ number of samples in group $g$;
Assign all samples belonging to test groups to the test set;
Assign all remaining samples to the training set;

**Table S1: Node Features attached to each residue node of PncA graphs.** Features were derived from either SBMLCore, STRIDE or a meta-predictor. Raw features were scaled based on the range of values in the train set.

| Node feature | Method |
|---|---|
| Amino acid volume | SBMLCore |
| Amino acid molecular weight | SBMLCore |
| Amino acid isoelectric point | SBMLCore |
| Amino acid hydropathy (Wimley–White) | SBMLCore |
| Amino acid hydropathy (Kyte–Doolittle) | SBMLCore |
| Num. hydrogen-bond acceptors | SBMLCore |
| Num. hydrogen-bond donors | SBMLCore |
| Num. side-chain rings | SBMLCore |
| Distance from PZA (drug) | SBMLCore |
| Distance from $Fe^{2+}$ | SBMLCore |
| Residue depth | SBMLCore |
| Solvent accessible surface area (SASA) | STRIDE |
| Phi ($\phi$) angle | STRIDE |
| Psi ($\psi$) angle | STRIDE |
| DeepDDG | Meta-predictor |
| SNAP2 | Meta-predictor |
| RaSP | Meta-predictor |
| MAPP | Meta-predictor |

**Table S2: Hyper-parameter tuning sweep parameter ranges.** Training was run for up to 3000 epochs with early stopping implemented where training would halt upon a plateau in test loss.

| Parameter | Value Range |
|---|---|
| Cutoff distance (Å) | [4.0, 18.0] |
| Dropout | [0.2, 0.8] |
| Edge weight method | *see Table 2* |
| Edge weight $\lambda$ (if exponential decay) | [0.1, 5.0] |
| Batch size | [32, 256] |
| Hidden channels | [32, 512] |
| Learning rate | $[10^{-7}, 10^{-2}]$ |
| Weight decay | $[10^{-7}, 10^{-2}]$ |

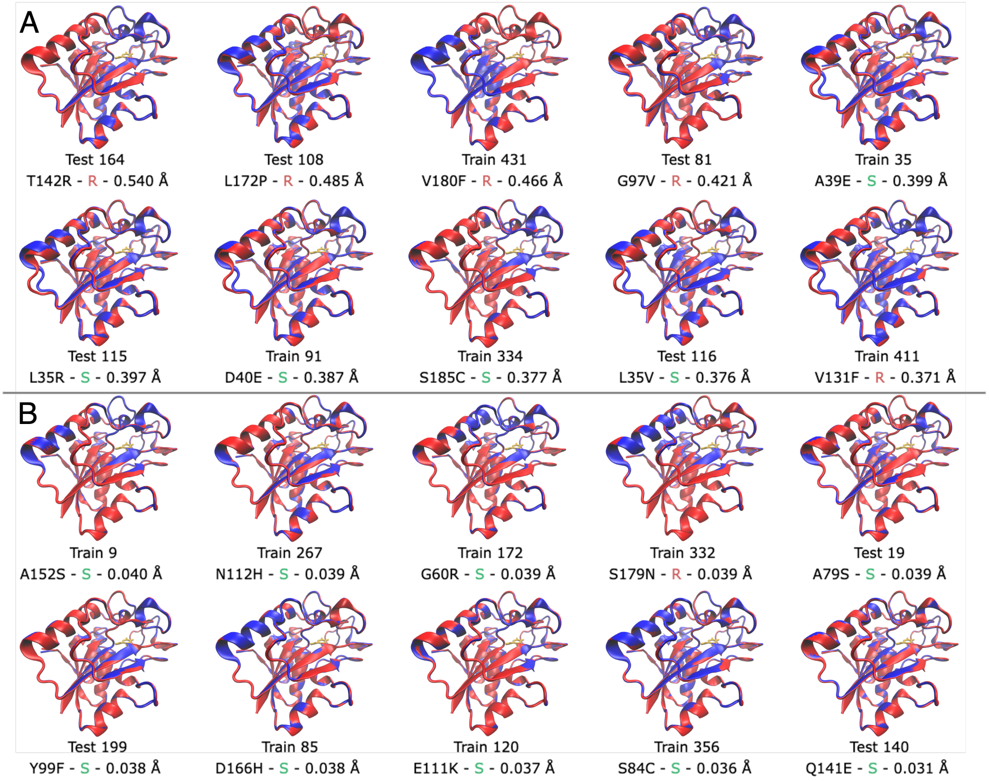


**Fig. S1: AlphaFold predicted structures aligned with AlphaFold prediction of wild-type.** Aligning the predicted structure (red) of the mutant variants with the wild-type PncA structure (blue) showed small but noticeable deviations in the structure. Each structure is labelled by its dataset (train or test) and index. The mutation, phenotype, and RMSD compared to the wild-type are all shown in each case. The ten structures with the highest **(A)** and lowest **(B)** RMSD are displayed.

**Table S3: Parameter count for trained GCN models.** Model size for each of the trained models is reported. The distinction between sizes is due to a different number of hidden channels found after hyper-parameter tuning for different datasets.

| **Model** | **Size (Number of Parameters)** |
|---|---:|
| GCN - Carter *et al.* split | 273,666 |
| GCN - Amino acid position split | 71,298 |
| GCN - Cluster split | 71,298 |

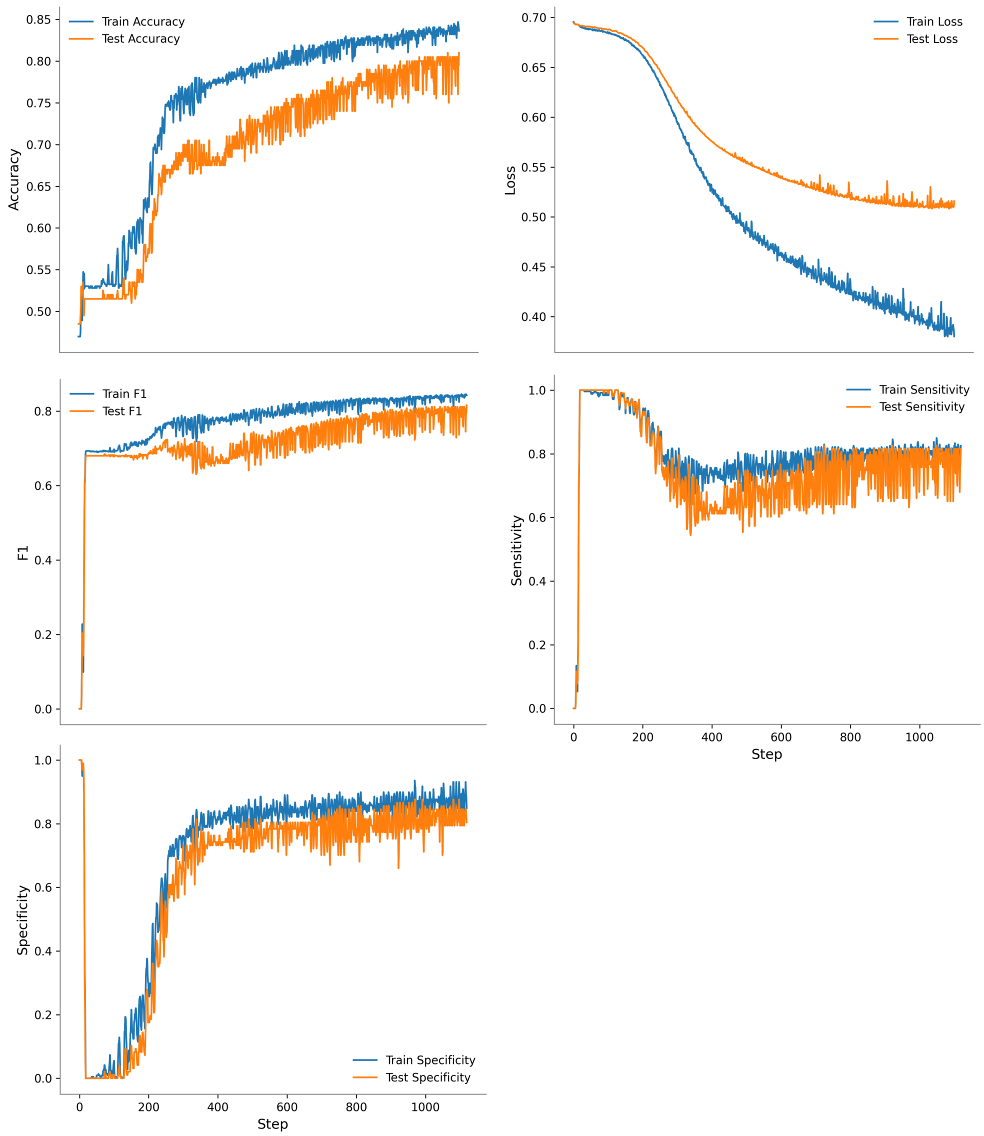


**Fig. S2: GCN model training curves.** The accuracy, loss, F1 score, sensitivity and specificity for the train test batches at each step. Training converged at step 1120 and the model at this step was used, which achieved the best performance.

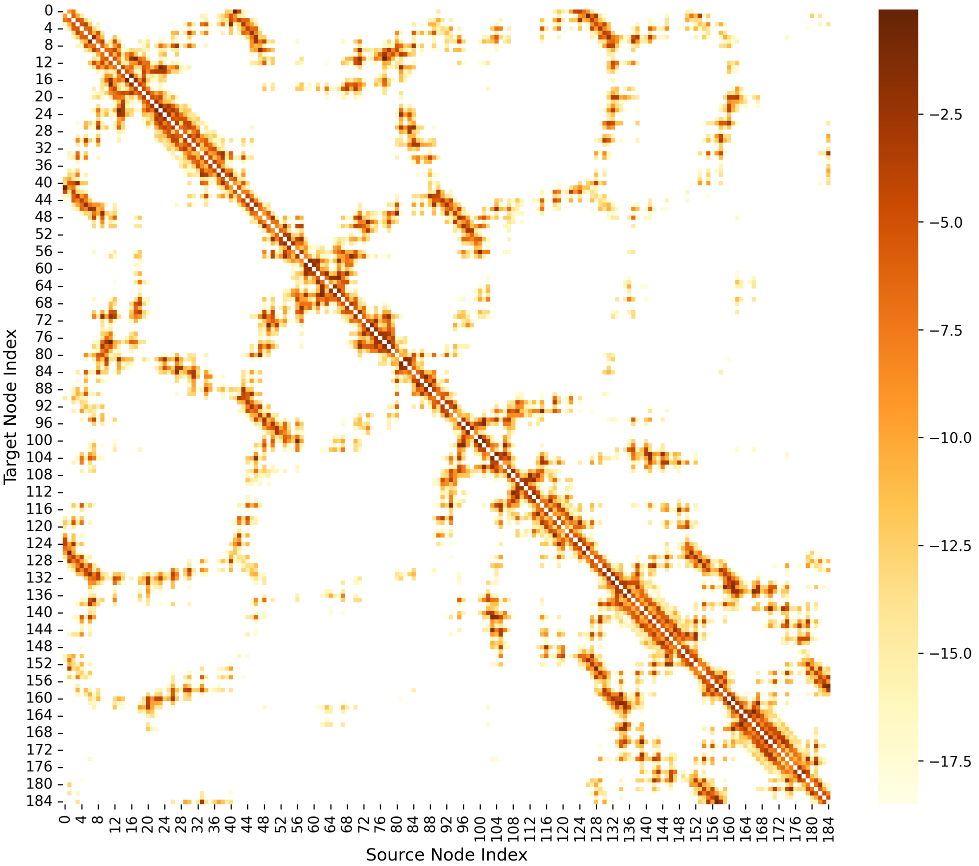


**Fig. S3: Edge weight heatmap for test set.** Heatmap shows the average edge weight across all PncA graphs in the test set for each given edge. Edge weight values are scaled logarithmically for visualisation purposes (as exponential edge weights were used). Edges that were not present in any PncA graph are shown as white in the heatmap.

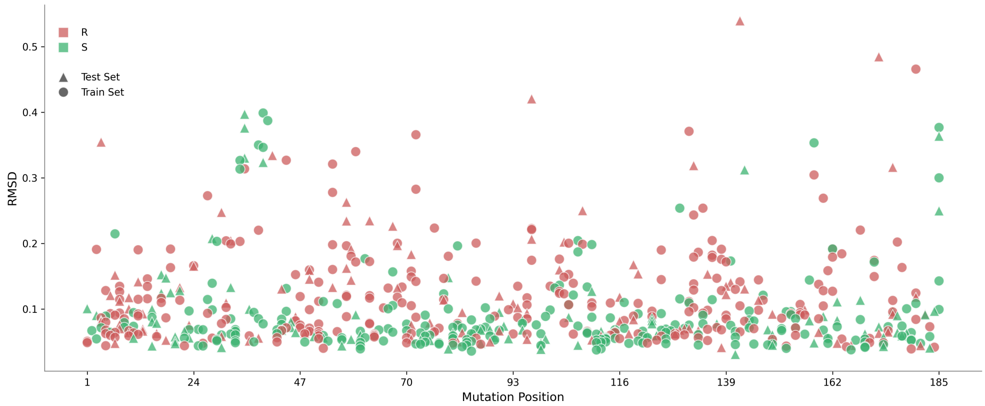


**Fig. S4: Scatterplot of RMSDs of AlphaFold2 structures by mutation position** Each structure is displayed as a triangle (test set) or a circle (train set), and coloured red (resistant) or green (susceptible).

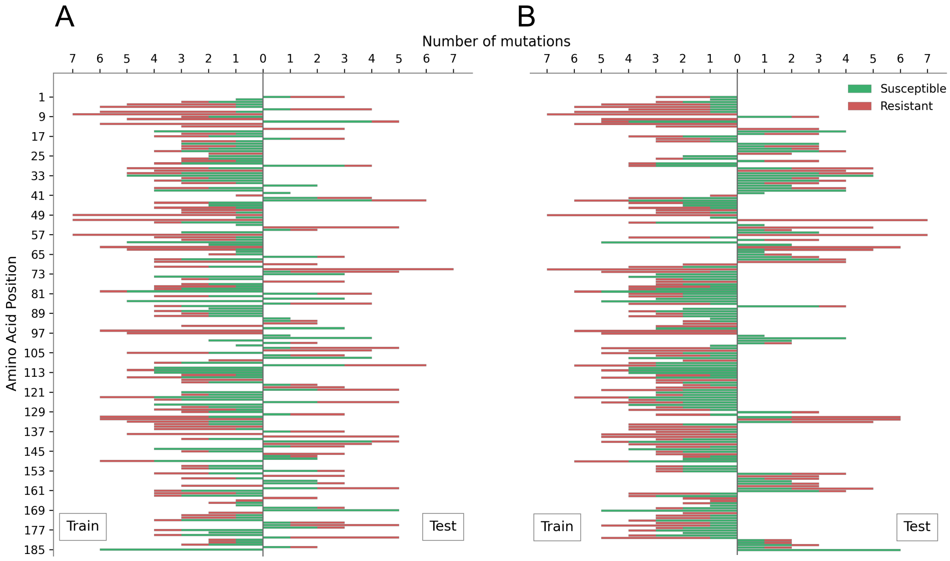


**Fig. S5: Distribution of mutations across the PncA protein sequence** Green and red indicate susceptible and resistant mutations respectively. Bars to the left and right indicate mutations found in samples in the train and test sets respectively, as per the following dataset splits. **(A)** Mutations in samples in the amino acid position split. All mutations at a given position are found entirely in either the train or test set and never in both. **(B)** Mutations in samples tructural cluster split.
